# Instantaneous change in hyphal diameter in basidiomycete fungi

**DOI:** 10.1101/2024.02.12.579893

**Authors:** Igor Mazheika, Oxana Voronko, Oxana Kolomiets, Olga Kamzolkina

**Author notes:** Corresponding author; Igor Mazheika.

## Abstract

Under certain conditions, fungi can rapidly change the size of their cells. For example, it is known that the cells of many yeast species under hyperosmosis instantly and reversibly shrink entirely, without plasmolysis, with a decrease in volume of up to 70%. There is evidence that filamentous fungi can also instantly change the diameter of their unspecialized hyphae. This property is fundamental but requires detailed study. In this large-scale study (involving more than 50,000 cells measured) using light microscopy, the ability of three unrelated basidiomycete species to rapidly change the diameter of their hyphae under various factors was analyzed. It was found that all three fungi respond similarly to moderate hyperosmotic shock or detergent treatment, shrinking by an average of 12–14% in diameter. However, inhibitors of actin assembly can cause either expansion or shrinkage of hyphae or have no effect on a fungus. These results, along with previously established features of the macroinvagination systems of the plasma membrane in basidiomycetes, are important for understanding the complex structural-protective physiological mechanisms responsible for the survival and continuous functioning of fungal cells in unstable environmental conditions.

## 1. INTRODUCTION

Fungi are extremely flexible organisms that quickly conform to their environment. One expression of fungal flexible is the ability to change the thickness of their hyphae (cell size in the case of yeast). Changes in hyphal diameters can occur over tens of min or hours during hyphal growth. For example, xylotrophic basidiomycetes, when inoculated from a rich nutrient medium to a medium without a nitrogen source, switch to in economical mode – the average thickness of *de novo* grown hyphae decreases sharply (Mazheika et al. 2020a). Another example can be found in the recent work showing that some fungi, for example, *Aspergillus nidulans* with an average hyphal diameter of 2–3 µm, easily grow through a 1 µm-width channel of a microfluidic device (Fukuda et al. 2021). One more example is the formation of giant yeast cells in *Cryptococcus neoformans* during pathogenesis to evade the host immune response and macrophage attacks (Zaragoza et al. 2010).

Changes in the diameters of fungal hyphae/cells can occur within tens of minutes, not necessarily accompanied by growth, due to a rapid regulatory response leading to changes in gene expression and physiological restructuring of the fungal cell. Thus, during moderate hyperosmotic shock in yeast and *Neurospora crassa* turgor pressure drops and the diameter of hyphae/cells quickly decreases (this rapid resizing is discussed in the next paragraph; Slaninova et al. 2000; Lew and Nasserifar 2009; Couttenier et al. 2022). Regulatory pathways are activated, primarily the CWI (Cell Wall Integrity) pathway, which also initiates the HOG (High Osmolarity Glycerol) pathway (de Oliveira et al. 2021). As a result, in less than half an hour, the synthesis of glycerol and other osmolytes begins. This, along with the regulation of ion uptake, allows the hyphae/cells to restore turgor pressure and their original size in about an hour (Lew and Nasserifar 2009; Ene et al. 2015).

Finally, the change in hyphae/cell diameter can occur instantly or near-instantly, within seconds to a maximum of several minutes, reversible and without loss of vitality. This is possible due to elasticity and the ability to rapidly remodel the cell wall, the tight adhesion of the plasma membrane to the cell wall, and the ability of the plasma membrane to quickly regulate the size of its working surface using macroinvaginations, along with other properties of the fungal cell (see Mazheika, Kamzolkina 2024 preprint). An example is the constricting rings of nematophagous fungi, which can significantly increase the diameter of the cells of the rings within a few seconds, thus trapping the nematode (Chen et al. 2022; Panstruga et al. 2023). Fundamentally important for mycology and this paper is another phenomenon: in the 1970s, it was observed that hyperosmotic shock could cause a sharp but reversible shrinkage of *Saccharomyces cerevisiae* and other yeast cells, with a volume loss of up to 70%. (Kopecká et al. 1973; Morris et al. 1986). Characteristic features of such cell shrinkage include significant thickening of the cell wall with remodeling of the glucan-chitin inner layer, absence or rarity of plasmolysis, formation and increase in the size of plasma membrane invaginations, and depolymerization of F-actin structures (Gray 1945; Kopecká et al. 1973; Morris et al. 1986; Slaninova et al. 2000; Ene et al. 2015; Palmer et al. 2015; Elhasi, Blomberg 2019; Lenardon et al. 2020; Couttenier et al. 2022; Gow, Lenardon 2023).

Filamentous fungi behave similarly to yeast under hyperosmotic conditions (Heath, Steinberg 1999; Lew et al. 2004; Lew, Nasserifar 2009; Lewis et al. 2009; Bitsikas et al. 2011; Walker, White 2017; Lew 2019). The differences lie in a less pronounced thickening of the wall and more developed systems of macroinvaginations of the plasma membrane (especially in xylotrophic basidiomycetes; Mazheika et al. 2022; Mazheika, Kamzolkina 2024 preprint). Unfortunately, few works are devoted to the rapid change in the diameter of hyphae with changes in osmotic and other environmental conditions. This process is described in detail in the work of Lew and Nasserifar (2009) on *N. crassa*. The authors showed that hyphae in 0.55 M sucrose shrink rapidly within about one minute, with the volume decreasing by approximately 40%, corresponding to a decrease in hyphal diameter of approximately 20– 25%. Chevalier and colleagues (2023, 2024) tested the effect of conditions with different osmolarities, showing a reduction in diameter of up to 35-40% in 1–1.5 M sorbitol on several species of filamentous ascomycetes and one zygomycete. However, they were working with apical hyphal cells, which differ from other hyphal cells in cell wall thickness and structure, turgor pressure, and possibly the cytoskeleton. Filamentous basidiomycetes have scarcely been studied in this context. Our recent paper (Mazheika et al. 2020b) demonstrated that 0.6 M sorbitol causes shrinkage of non-apical hyphae of *Rhizoctonia solani* J.G. Kühn by approximately 15%.

At the same time, the ability of fungi to instantly change the diameter of hyphae is an important part of the universal and complex mechanism of fungal adjustment to rapid changes in external conditions, both minor and stressful. This structural-protective mechanism is especially interesting in xylotrophic, mycorrhizal, and other basidiomycetes, which are known for their complexity of organization and physiology at both the colony and intrahyphal levels (Brushaber and Jenkins 1971; Ashford 1998; Weber et al. 1998; Watkinson et al. 2005; Zhuang et al. 2009; Schmieder et al. 2019; Sheldrake 2020; Mazheika et al. 2020a, 2022). The goal of this work is to assess the ability of three species of basidiomycetes from different systematic and ecological groups to change the thickness of their hyphae in response to various factors, including osmotically active and membrane-permeabilizing agents, and inhibitors of F-actin assembly. The study is based on a light microscopic quantitative and qualitative analysis, using a large number of fungal cells (several tens of thousands). The obtained results, along with those from our previous studies on the systems of macroinvaginations of the plasma membrane and the structural-physiological model of fungi (the curtain model; Mazheika et al. 2020b, 2022; Mazheika, Kamzolkina 2024 preprint), will represent another step towards a holistic understanding of fungal cell physiology and the complex mechanisms of fine adjustment to unstable environmental conditions.

## 2. MATERIAL AND METHODS

### 2.1. Strains and culture conditions

Wild-type strains of three species of basidiomycetes were used: *Stereum hirsutum* (Willd.) Pers. (Mazheika et al. 2020a); *Rhizoctonia solani* (Kamzolkina et al. 2017; Mazheika et al. 2020b); *Coprinus comatus* (O.F. Müll.) Pers. (Mazheika et al. 2020a). Mycelium was grown and stored on malt agar with 1.5% agar (Panreac, Spain; Mazheika et al. 2020a). To prepare microscopic samples, the material was grown on the same malt agar in 9 cm Petri dishes, but the nutrient medium was covered with a cellophane disk (Mazheika et al. 2022). Colonies 4.5 d old were most often used (colonies should not reach the edge of the dish: *R. solani* and *S. hirsutum* were inoculated to the edge of the dish, *C. comatus* in the center).

### 2.2. Preparation of light microscopic samples, microscopy and photography

To prepare the microscopic slides, a piece of cellophane (approximately 7 ˟ 10 mm) with a fragment of the colony was cut out with a blade to include the marginal mycelium and a couple of millimeters of mycelium with secondary growth. In most cases, the slide was mounted with 30 µL of malt medium (the same as the growth medium but without agar), either without any additives or with various additives. The corresponding amount of solvent for the active component (water, buffer, or DMSO – dimethyl sulfoxide; Sigma-Aldrich, Missouri) was added to the malt medium in control slides.

Experiments with control scatter to determine the biological and technical errors were conducted as follows: three samples were taken from one Petri dish with a fungal colony. The samples were cut equidistant from each other and not near the edge of the dish (if the inoculum was placed at the edge). The slides were mounted with malt medium.

To create hyperosmotic conditions, slides were mounted with malt medium with added sorbitol (Sigma-Aldrich) at a final concentration of 0.6 M. To test the effect of the detergent, slides were mounted with a solution in which Triton X-100 (10% water stock; Sigma-Aldrich) was added to the malt medium or in malt medium with 0.6 M sorbitol to a final concentration of 1% v/v. To study the effect of proteolysis, a 20 mg/mL stock of Proteinase K (Medgen, Russia, 30ea) in 0.1 M Tris-HCl pH 8 with 1.5 mM CaCl_2_ (Chimmed, Russia) and 50% v/v glycerol (Chimmed) was prepared. The stock was added to the malt mounting medium, adjusted to pH 8 with KOH, to a final concentration of 1 mg/mL. An additional control was also used: instead of proteinase K, an equimolar amount of BSA (bovine serum albumin; AppliChem, Germany) was added at a final concentration of 2.3 mg/mL to the mounting medium. To create low osmolarity conditions, samples were mounted with distilled water instead of malt medium. The effect of F-actin disassembly on hyphal diameter was tested using the actin assembly inhibitors latrunculin A (LatA; Santa Cruz Biotechnology, Texas) and cytochalasin D (CytD; Enzo, Switzerland), which were added to malt medium from DMSO stocks at final concentrations of 50 to 200 µM.

In the series of experiments with sample preincubation in liquid medium, the usual malt control and the sample with 0.6 M sorbitol were first cut out from the Petri dish and analyzed. Then, the same cellophane disc with the fungal colony was transferred to a fresh Petri dish without malt agar, and 15 mL of malt medium was poured. After 45 min of incubation in the dark at room temperature, the third sample was cut out, and the slide was mounted with the same malt medium. After 75 min, the fourth sample was cut out and mounted with malt medium containing sorbitol.

The influence of mechanical stress was studied as follows: the sample was placed on a slide, mounted with malt medium, covered with a coverslip, and then lightly tapped on the coverslip with a plastic tube. The samples were immediately subjected to microscopic analysis.

Immediately after preparing the slide, it was placed under a Zeiss Imager M1 microscope (Germany) equipped with a Hamamatsu 1394 ORCA-ERA camera (Japan). Samples were photographed in light transmission mode using ˟20 lens and a constant shutter speed of 100 ms, with adjusted light intensity. Photographs were taken over 25 min, divided into two observation intervals: 0–10 min and 10–25 min. Approximately 40 to 80 fields of view were photographed during each interval. The fields of view were not chosen randomly: photographs were taken primarily to avoid the colony’s edge with numerous apical hyphae, also the basal zones with a dense plexus of hyphae were not photographed. They were choosen also so that in the field of view there were septa or septa with clamp connections. For each mycelium treatment option and for the control scatter, at least three biological replicates were set (a repeat was considered to be a sample taken from a separate Petri dish with mycelium, grown in most cases in different batches).

In the dynamic experiments, the sample was mounted with malt medium containing 0.6 M sorbitol. The slide was quickly installed in the microscope, a suitable field of view was found, and photographs were taken at 60-second intervals. Since changes and loss of focus could occur over time, the focus was carefully adjusted before each photograph, using predetermined marker hyphae.

### 2.3. Processing of results, computer programs used

IMAGEJ 1.54d (Schneider et al. 2012) was used to measure hyphal diameters using the Straight tool and the Measure function. The hyphae located in a specific focal section were measured. The focus during shooting was adjusted to minimize the thickness of the cell wall while maximizing contrast, ensuring the cell diameter was at its largest. Cells in other focal sections were ignored. Measurements were taken on the outer clear contrast boundaries of the cell wall. The following rules guided mesure site selection: apical hyphae were not measured (the tip should not be visible in the field of view), and severely uneven or swollen hyphae were excluded. Only hyphae were measured that have septa in *R. solani*; complex clamp connections with at least two connections in *S. hirsutum* (usually, the hyphae with one connection in this species are very thin); and regular clamp connections in *C. comatus* (in this species, all clamp connections have one connection; see FIG. 1A–C). Measurements were taken at a distance from 20 µm to several hundred µm from the septum or clamp connection. If the hypha has different thickness along its length, it was measured at the widest point.

**Figure 1.**
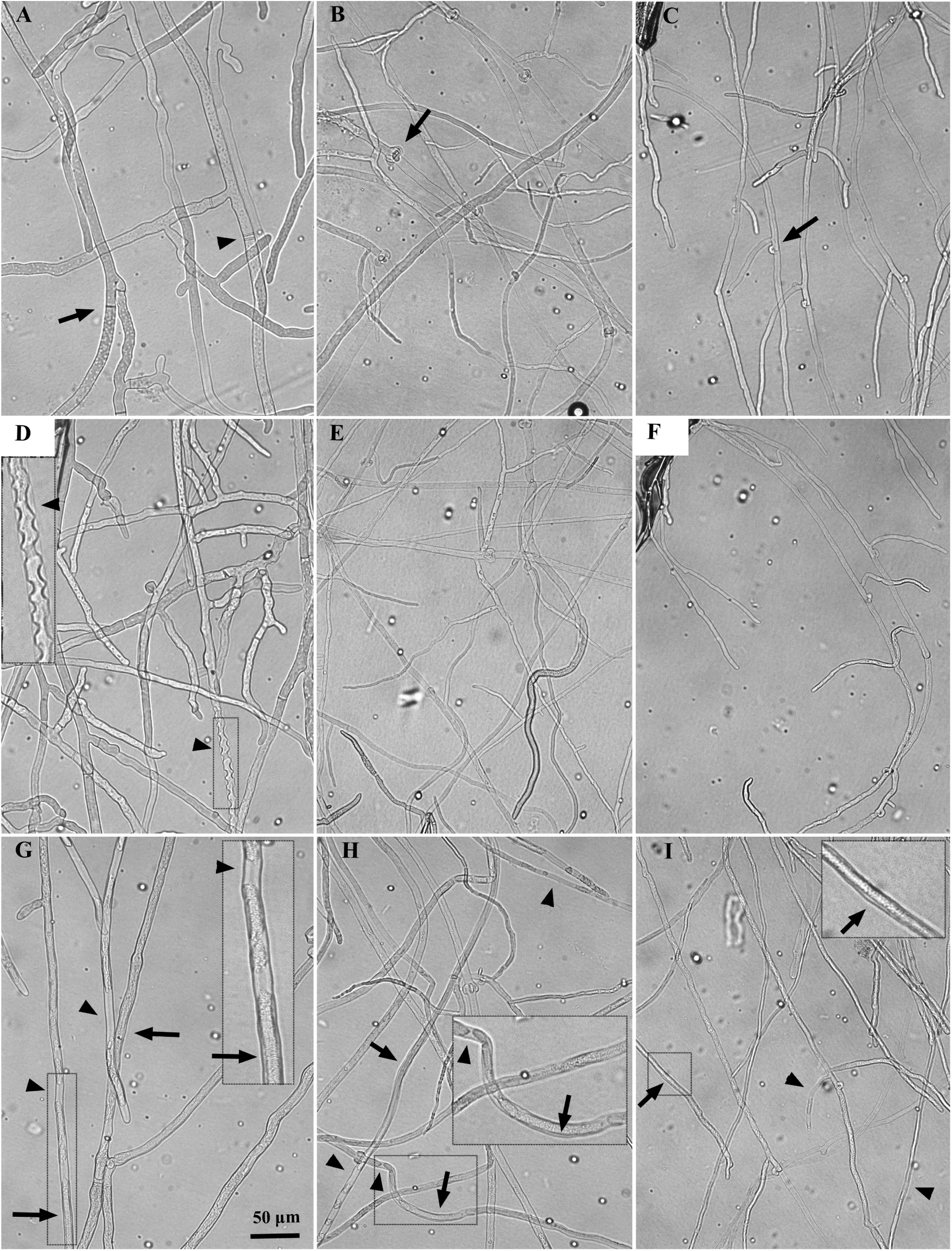
Bright-field microscopic photographs of the mycelium of the three studied fungi under different processing conditions. A–C: control mycelium (mounted with malt medium) of *R. solani* (A), *S. hirsutum* (B) and *C. comatus* (C). The arrows indicate the septum without a clamp connection in *R. solani*, the septum with a complex clamp connection in *S. hirsutum*, and the septum with a regular clamp connection in *C. comatus*. The arrowhead shows a double septum in *R. solani*. D–F: microscopic slides mounted with medium containing 0.6 M sorbitol. Even without measurements, it is clear that the hyphae, on average, become narrower relative to the control. At the same time, *S. hirsutum* (E) and *C. comatus* (F) exhibit practically no other morphological changes or plasmolysis. In *R. solani* (D), concave plasmolysis is observed in some cells (arrowhead) within the first 10 min after slide preparation. G–I: mycelium treatment with 1% Triton X-100. In all three fungi, *R. solani* (G), *S. hirsutum* (H), and *C. comatus* (I), shrinkage of the hyphae and convex plasmolysis (arrows) are observed, with splitting of the protoplast into parts (arrowhead) in some cells. The dotted frames indicate fragments of photographs (D and G–I), which are enlarged for greater detail and placed within the same photograph.

Data were then transferred to EXEL (Microsoft, Washington), ORIGIN PRO 7.0 (OriginLab, England) and STATISTICA 6 (StatSoft, Oklahoma). Scatter plots were generated in ORIGIN PRO. All statistical calculations were performed in STATISTICA 6. Paired samples were compared using the Mann-Whitney U test. Differences in the average diameters of hyphae (D, in µm) between the control and the sample were assessed as a percentage using the formula _Δ_D = 100 − (D_1_ × 100)/D_2_, where D_1_ and D_2_ are the average diameters of the hyphae in the control and sample, with the smaller number in the numerator and the larger one in the denominator. If the average diameter of the sample is greater than that of the control (hyphal expansion), a “−” sign was placed in front of the _Δ_D value. _Δ_D values were rounded to one decimal place. Similarly, _Δ_D was used to compare two observation time intervals, 0–10 min and 10–25 min, within one control/sample. Where possible, the mean change in diameters (mean _Δ_D) and standard deviation (SD) between independent experiments were calculated.

In experiments establishing the control scatter of hyphal diameters, _Δ_D was determined between the two extreme values of D among three samples from the same Petri dish. At least three independent experiments were conducted, and the sum of the mean _Δ_D and SD was calculated (_Δ_D + SD) and rounded to the nearest higher half-unit. In this study, any _Δ_D was considered significant if it was greater than this value in modulus and *P* (U-test) < 0.05. If _Δ_D was less than this value, it indicated no change in the thickness of the hyphae even at *P* (U-test) < 0.01.

When processing the data from the dynamic experiments, three to six non-apical hyphae were selected in each sample. Three points, distant from each other, were taken on each hypha, and their diameters were measured in each frame of the time-lapse video. For each measurement point, the Spearman correlation coefficient (r_s_) was calculated between hyphal diameter and observation time. It was assumed that the hypha at a given point successively expands or shrinks at r_s_ ≥ 0.4 and r_s_ ≤ −0.4, *P* < 0.01, respectively.

Images and videos were generated and processed in PHOTOSHOP CS (Adobe Systems, California) and IMAGEJ 1.54d.

## 3. RESULTS

### 3.1. Results underlying all other findings obtained in the study

During the work, approximately 55 000 cell diameters in the three studied fungal species were measured (20 146 for *S. hirsutum*; 17 832 for *R. solani*; 17 072 for *C. comatus*). The average diameter of control hyphae decreases in the order: *R. solani*, *S. hirsutum* and *C. comatus*; and is 9.7± 0.4, 7.9 ± 0.5 and 5.6 ± 0.2 µm, respectively (after the sign ± SD is indicated hereinafter). The growth rate of the mycelium also decreases in this order (data not shown). It is important to note the following underlying results: (i) a sharp change in hyphal diameters when mounting slides with hyperosmotic and some other media occurs very quickly (within the first few minutes or faster). With our methodological approaches, such rapid expansion or shrinkage of hyphae is difficult to record directly due to the time interval between preparing the specimen and placing it under the microscope (about 1.5–2 min). Therefore, conclusions about rapid shrinkage/expansion are made indirectly based on: first, the presence of significant differences between the average diameters of the samples and their controls (−4.5% > _Δ_D > 4.5%). However, such differences do not indicate a mandatory rapid change in the diameters of the hyphae in the sample; perhaps the hyphae gradually shrank or expanded during the 25 min of observation. Therefore, D of the two observation intervals of 0–10 and 10–25 min was also compared. For example, in 21 samples with sorbitol (samples of all three fungi combined), only in one sample did D (0–10 min) ≠ D (10–25 min), *P* (U-test) < 0.05, indicating a rapid change in D in the first minutes and then its preservation within 25 min in most sorbitol samples. Second, the dynamic experiments, where there is no general tendency for a gradual change in hyphal diameters over the course of an hour after shrinkage in sorbitol, also confirm that the change in diameter occurs within the first seconds to minutes after slide preparation. (ii) Once shrunk (or expanded), the hyphae generally remain in this state without returning to their original size, at least for an hour, as observed in dynamic experiments. However, as indicated below, in a small number of samples, there is a tendency for some gradual change in D over the 25-min observation interval, but these changes are not necessarily aimed at restoring the original hyphae sizes. (iii) Hyphae shrink or expand as a whole, together with the cell wall. Plasmolysis (detachment of the protoplast from the cell wall) rarely occurs, but there are some exceptions: *R. solani* exhibits fast reversible concave plasmolysis within the first minutes in the hyperphase (FIG. 1D); *S. hirsutum* shows a similar effect after one hour of preincubation in liquid medium; and treatment with a detergent induces irreversible convex plasmolysis in all fungi (FIG. 1G–I). However, in all cases, even under the action of a powerful detergent, plasmolysis does not occur in all hyphal cells. (iv) A statistically significant _Δ_D change reflects the general trend in changes in hyphal diameters in the sample. However, the behavior of individual hyphae and even neighboring sections of the same hyphae may not coincide with the general trend. Indicative in this case are dynamic experiments in which the hypha can gradually narrow at one measurement point, and expand at a neighboring point. (v) The native D of the hyphae before placing the mycelium on a microscopic slide is not known: when mounting the slide, even with the control malt medium, the D of the mycelium may change rapidly and undetectably relative to the original mycelium in the Petri dish.

### 3.2. The influence of various factors on changes in the diameter of hyphae in three studied species of fungi

For further work, it was necessary to establish the value of the control scatter to accept the minimum value of _Δ_D, below which changes in the size of the hyphae can be neglected. This scatter arises due to both technical measurement errors and biological reasons – specifically, the heterogeneity of the growth pattern within one fungal colony. It turned out that, despite different growth rates and different average thickness of hyphae in the three species of fungi, their control spread of D is very close (FIG. 2A). For all three fungi, the control mean _Δ_D + SD is 4.0–4.1%, and the maximum control _Δ_D is 3.9–4.3%. Accordingly, we accepted _Δ_D = 4.5% for all three fungi as a threshold below which (in absolute value; 4.5 is not included), even at *P* (U-test) < 0.01, the change in hyphal diameter should be disregarded. For example, if in the control the average diameter of the hyphae was 10 µm, and in the sample it was 9.6 µm (_Δ_D = 4%), the difference was disregarded for any *P*; if the sample had 9.5 µm (_Δ_D = 5%), the difference was considered significant only if *P* (U-test) < 0.05.

**Figure 2.**
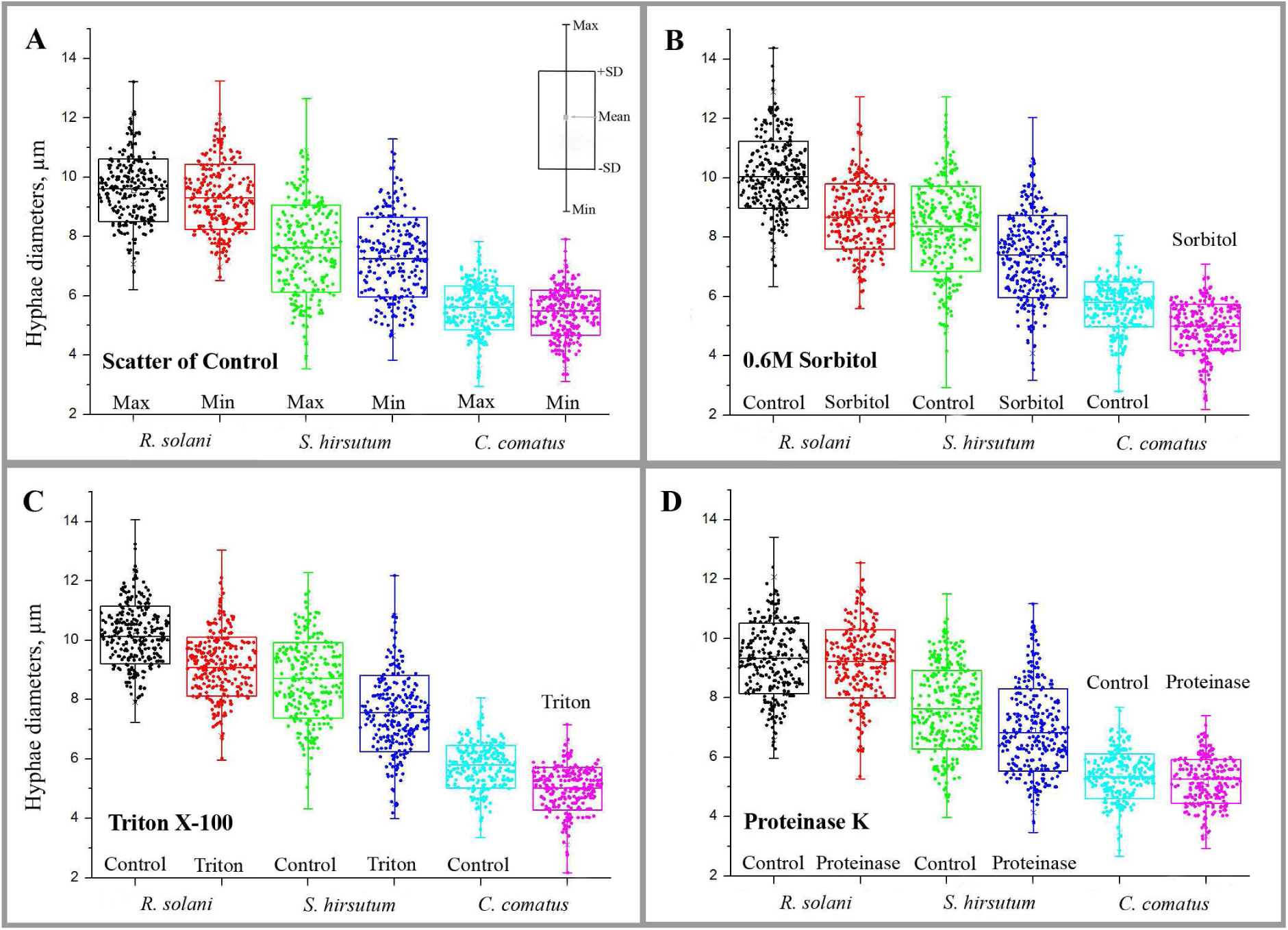
Combined diagrams illustrating the influence of three factors (0.6 M sorbitol, 1% Triton X-100, and 1 mg/mL Proteinase K) on the instantaneous change in hyphal diameters in three studied fungi. For comparison, a diagram of the internal scatter of diameters in the control (A) is included. Each column of the chart combines samples from 3–9 experiments. Each sample is the sum of samples from two time intervals: 0–10 min and 10–25 min. Each factor is represented by two columns: control and sample (Sorbitol, Triton, or Proteinase). In panel A, the column labeled Max combines samples with the largest average hyphal diameters in their experiments, while Min combines those with the smallest. The diagrams show that mounting microscopic slides with 0.6 M sorbitol (B) or Triton X-100 (C) leads to pronounced shrinkage of hyphae relative to the control. *R. solani* and *S. hirsutum* each have one outlier experiment excluded in panel B. The effect of Proteinase K (D) is not as pronounced; only *S. hirsutum* shows a general tendency to shrink hyphae.

#### 3.2.1. Influence of conditions with high and low osmotic medium

Mounting the slides with the hypertonic medium based on 0.6 M sorbitol in all three fungi causes significant shrinkage of the hyphae (FIG. 2B). In only one out of seven experiments, *R. solani* had _Δ_D < 4.5% (indicating no general shrinkage of the hyphae). The degree of shrinkage among the three fungi is similar: on average, *S. hirsutum* shrinks the least (mean _Δ_D = 11.6 ± 2.6%; nine independent experiments; min _Δ_D = 8.2%, max _Δ_D = 16.5%), and *R. solani* shrinks the most (mean _Δ_D = 14.3 ± 2.4%; six independent experiments; min _Δ_D = 10.5%, max _Δ_D = 17.5%; excluding one experiment without shrinkage). *C. comatus* has mean _Δ_D = 11.9 ± 4.0%; five independent experiments; min _Δ_D = 5.6%, max _Δ_D = 15.4%. For all _Δ_D *P* (U-test) < 0.01. In *S. hirsutum* and *C. comatus*, hyphae in sorbitol shrink without plasmolysis (concave plasmolysis was observed in only one out of nine experiments in *S. hirsutum*). In *R. solani*, the situation is different: concave plasmolysis occurs in some cells within the first 8−10 min after mounting a slide (the incidence varies between experiments), which then disappears (FIG. 1D; SUPPLEMENTARY VIDEO).

When comparing D at observation time intervals of 0−10 min and 10−25 min in the experiments with sorbitol, the results are as follows. In all five experiments with *C. comatus* and in seven experiments with *R. solani*, both in the samples and controls, there was no difference between the intervals of 0−10 min and 10−25 min. In *S. hirsutum*, only in one experiment out of nine did the sorbitol sample have D (0–10 min) > D (10–25 min), indicating that after the sharp shrinkage, the hyphae continued to gradually shrink further (_Δ_D = 7.2%). However, in another experiment, the control had the same result.

It turned out that preincubating *S. hirsutum* mycelium in malt medium before mounting the slides affects the behavior of the mycelium in microscopic samples. In two of three independent experiments, preincubation in malt medium caused slight hyphal expansion (_Δ_D = −5.5% and −7.5%) when comparing the controls before and after preincubation. In one experiment, preincubation did not affect the average thickness of the hyphae but did influence the reaction to sorbitol: this mycelium, after preincubation, did not shrink when mounted with sorbitol-containing medium. In the first two experiments, mycelium shrinkage in sorbitol occurred but was less than when treated with sorbitol without preincubation: _Δ_D = 7.7% and 7.0%. The mycelium shrank approximately as much as it expanded after preincubation in malt medium. However, FIG. 3A shows that such shrinkage is accompanied by an increase in the spread (SD) of diameters. Compared to the initial mycelium, the mycelium that has undergone preincubation and sorbitol treatment has a certain number of both very narrow and very wide hyphae, although the average diameter of the hyphae remained virtually unchanged. In addition, preincubation has another important effect. In the two experiments where hyphal expansion and shrinkage occurred, concave plasmolysis was observed in many cells of the preincubated mycelium within the first 5−10 min after mounting slides with sorbitol-containing medium. In this case, the behavior of the *S. hirsutum* mycelium resembles the reaction of *R. solani* mycelium to sorbitol without preincubation. In the experiment where expansion/shrinkage did not occur, there was no plasmolysis.

**Figure 3.**
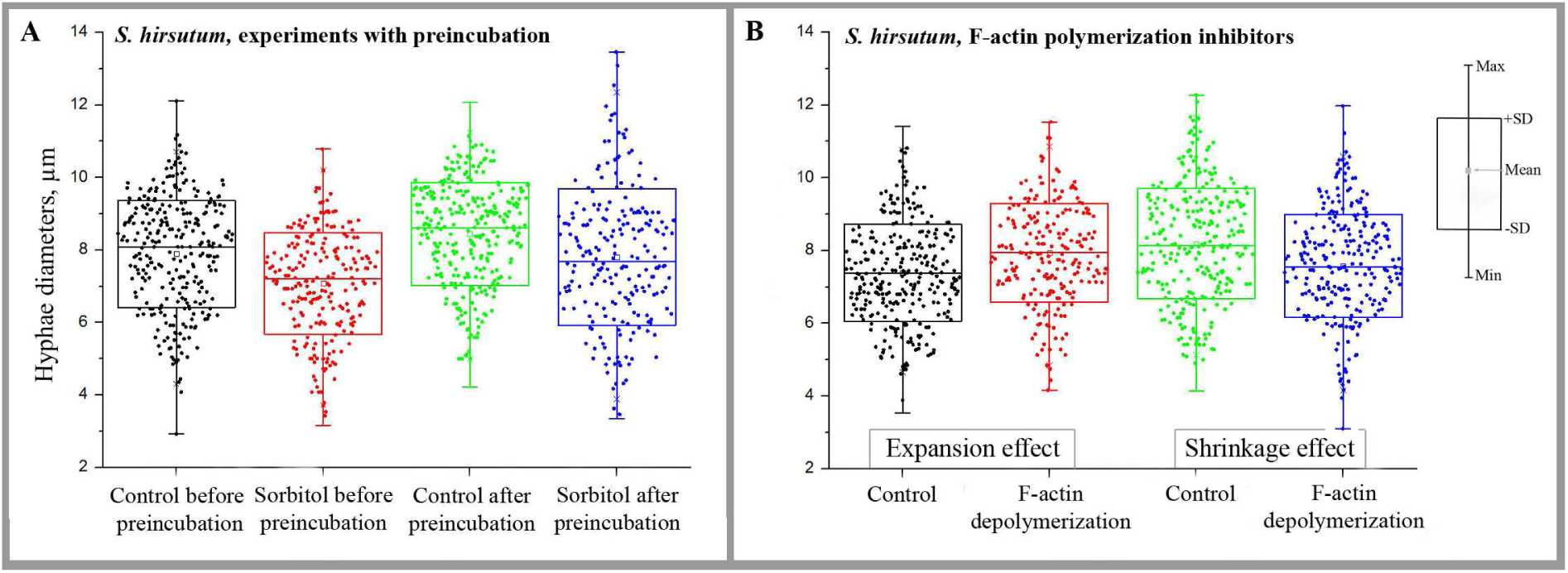
Combined diagrams illustrating the influence of mycelium preincubation in malt medium (A) and inhibitors of F-actin assembly (B) on the diameter of *S. hirsutum* hyphae. In panel A, samples from two experiments were combined. It is observed that preincubation of the mycelium in the liquid medium results in a slight expansion of the hyphae (Control after preincubation) compared to the usual control (Control before preincubation). Mounting samples, both without preincubation (Sorbitol before preincubation) and with preincubation (Sorbitol after preincubation), in sorbitol leads to shrinkage of the mycelium. However, the preincubated mycelium shrinks less on average, and its sampling spread is greater than that of the controls. Diagram B combines four experiments with different concentrations of LatA (there is no correlation between the dose of the inhibitor and the response to it) and 100 µM CytD. Diagram columns combining the two samples with their controls, showing an increase in average hyphal diameter after treatment with actin inhibitors, are labeled “Expansion effect.” Accordingly, the “Shrinkage effect” columns contain two samples that shrank after treatment with the inhibitors relative to their controls. It can be seen that the expanding mycelium has the control being on average narrower in diameter compared to the control of the shrinking mycelium.

Distilled water, as a mounting medium with low osmolarity, has little effect on changes in hyphal diameters. It has almost no effect on *C. comatus* and *S. hirsutum* (in only one out of three experiments, the hyphae of *S. hirsutum* shrank by 4.9%). In *R. solani*, in one experiment out of three, it caused the expansion of hyphae by 7%. Moreover, in all experiments with water, there is no difference between D (0–10 min) and D (10–25 min), indicating that water does not cause an immediate or gradual change in D.

#### 3.2.2. Effects of plasma membrane permeabilizing agents

Triton X-100 added to the mounting medium produces an effect quantitatively similar to that of sorbitol. In all three fungi, the hyphae shrink by about 11–13% (FIG. 2C). There is no statistical difference between the shrinkage in sorbitol and Triton X-100. There is also no statistical difference between samples with Triton X-100 and those with the simultaneous addition of Triton X-100 and sorbitol. More detailed information for *S. hirsutum*: mean _Δ_D = 12.5 ± 4.6%; three independent experiments; min _Δ_D = 7.2%, max _Δ_D = 15.5%. For *R. solani*: mean _Δ_D = 10.8 ± 0.5%; three independent experiments; min _Δ_D = 10.3%, max _Δ_D = 11.3%. For *C. comatus*: mean _Δ_D = 13.1 ± 1.4%; three independent experiments; min _Δ_D = 12.3%, max _Δ_D = 14.8%. For all _Δ_D *P* (U-test) < 0.01. However, unlike sorbitol, Triton X-100 induces strong convex plasmolysis in some cells with protoplast splitting in all three fungi (FIG. 1H–I). In *C. comatus*, such plasmolysis occurs less frequently than in the other two fungi. In *R. solani*, convex plasmolysis occurs more frequently in the apical cells than in the other cells.

In all slides with the addition of Triton X-100 and Triton X-100 with sorbitol, there is no difference in the hyphal diameters between the observation intervals of 0–10 min and 10– 25 min in *S. hirsutum* and *R. solani*. In *C. comatus*, in all three independent experiments, in slides with simultaneously added Triton X-100 and sorbitol, D (0–10 min) ≠ D (10–25 min), *P* (U-test) < 0.01. However, of these three experiments, in two there is a weak gradual expansion of hyphae _Δ_D = −5.4% and −5.8%, and in the third, a very weak shrinkage _Δ_D = 4.6%. The effect of Proteinase K on D of the studied fungi is not as strong and uniform as the effects of sorbitol and Triton X-100 (FIG. 2D). Proteinase K has the greatest effect on *S. hirsutum*: out of three experiments, the enzyme had no effect in one, caused a slight shrinkage of 5.2% in another, and resulted in significant shrinkage of 15.9% in the third. In *C. comatus*, Proteinase K caused hyphal shrinkage by 8.8% in only one out of three experiments. Interestingly, in the same slide, D (0–10 min) > D (10–25 min), with _Δ_D = 6.9%. In other words, the hyphal shrinkage occurred gradually rather than instantly. Proteinase K had almost no effect on the mycelium of *R. solani*; only in one experiment was there a slight shrinkage of 5%. There was no difference between the observation time intervals of 0–10 min and 10–25 min for either *R. solani* or *S. hirsutum*.

Despite Proteinase K having minimal effect on the average diameter of *R. solani* hyphae, concave plasmolysis can be observed in some cells within the 0–10 min interval. Additionally, in the first 10 min, single cells with concave plasmolysis and individual thin hyphae with convex plasmolysis are noted in *C. comatus* during the experiment where hyphal shrinkage occurs in Proteinase K. There is no plasmolysis in *S. hirsutum*.

BSA added to the mounting medium instead of Proteinase K in equimolar amounts had no effect on the hyphal diameter of *S. hirsutum* and *R. solani*. However, in the same experiment where Proteinase K caused hyphal shrinkage in *C. comatus*, BSA also resulted in a shrinkage of 7.2%. In another experiment with *C. comatus*, BSA caused a hyphal expansion of 6.5%.

#### 3.2.3. F-actin disruption and mechanical stress as influencing factors

Inhibitors of F-actin assembly, LatA and CytD, have an ambiguous effect on the fungi studied. They do not affect the diameter of the hyphae of *R. solani* and *C. comatus*. In only one out of four experiments, 50 µM LatA causes a gradual expansion of hyphae in *R. solani*: D (0–10 min) < D (10–25 min), _Δ_D = –7.8%. However, there is still no overall difference between the control and the sample in this experiment.

The action of LatA and CitD on *S. hirsutum* is as follows: (i) the effect of the inhibitors on D is dose-independent, with r_s_ = 0.07 for the dependence of _Δ_D on the concentration of the inhibitor. (ii) In five of nine experiments, the inhibitors did not affect the diameters of hyphae. In two experiments, they caused expansion of hyphae by 6.0% (100 µM CytD) and 6.6% (200 µM LatA), and in two experiments, shrinkage by 7.2% (50 and 100 µM LatA; FIG. 3B). Only in one slide with 50 µM LatA (no overall effect on hyphal diameter), there is a slight gradual shrinkage of hyphae: D (0–10 min) > D (10–25 min), _Δ_D = 4.8%.

An additional variant of the experiment was conducted on *S. hirsutum* involving mechanical stress. Tapping the cover glass of the slide caused slight shrinkage of the hyphae compared to the control: in one out of three experiments, there was no change in hyphal diameter, while in two experiments, there was shrinkage by 4.8% and 6.9%. There was no difference in observation intervals of 0–10 and 10–25 min. However, damage and crowding of hyphae observed in the slides could have resulted in unreliable hyphal diameter measurements.

### 3.3. Behavior of the mycelium of the studied fungi in dynamic experiments

All the experiments described above deal with average values among numerous quasi-random hyphae. The dynamic experiments have a different approach. Changes in the diameters of individual hyphae were analyzed at fixed measuring points within an hour after preparing the microscopic specimen with sorbitol. The following results were obtained (FIG. 4): (i) none of the 25 individual hyphae of three studied species of fungi show a general tendency to expand or shrink after exposure to sorbitol (|r_s_| ≥ 0.40) – if a gradual change in diameter occurs, it happens only at one or two points out of three selected on each hypha, but never at all three. (ii) Only 17–27%, depending on the species of fungus, of the points selected on the hyphae show a gradual change in diameter. At the remaining points, either no changes occur within an hour, or they fluctuate back and forth, making them unsuitable for description by linear Spearman correlation. (iii) There is a difference in dynamic experiments between the three fungal species: *S. hirsutum* has the most hyphal sections with a tendency to gradually change in diameter (about 27%; FIG. 4A–B), with half tending to expand and half to shrink. In *R. solani*, only about 17% of the hyphal sections tend to change, all becoming narrower (FIG. 4C). In *C. comatus*, 19% of points are variable; however, all tend to expand (FIG. 4D).

**Figure 4.**
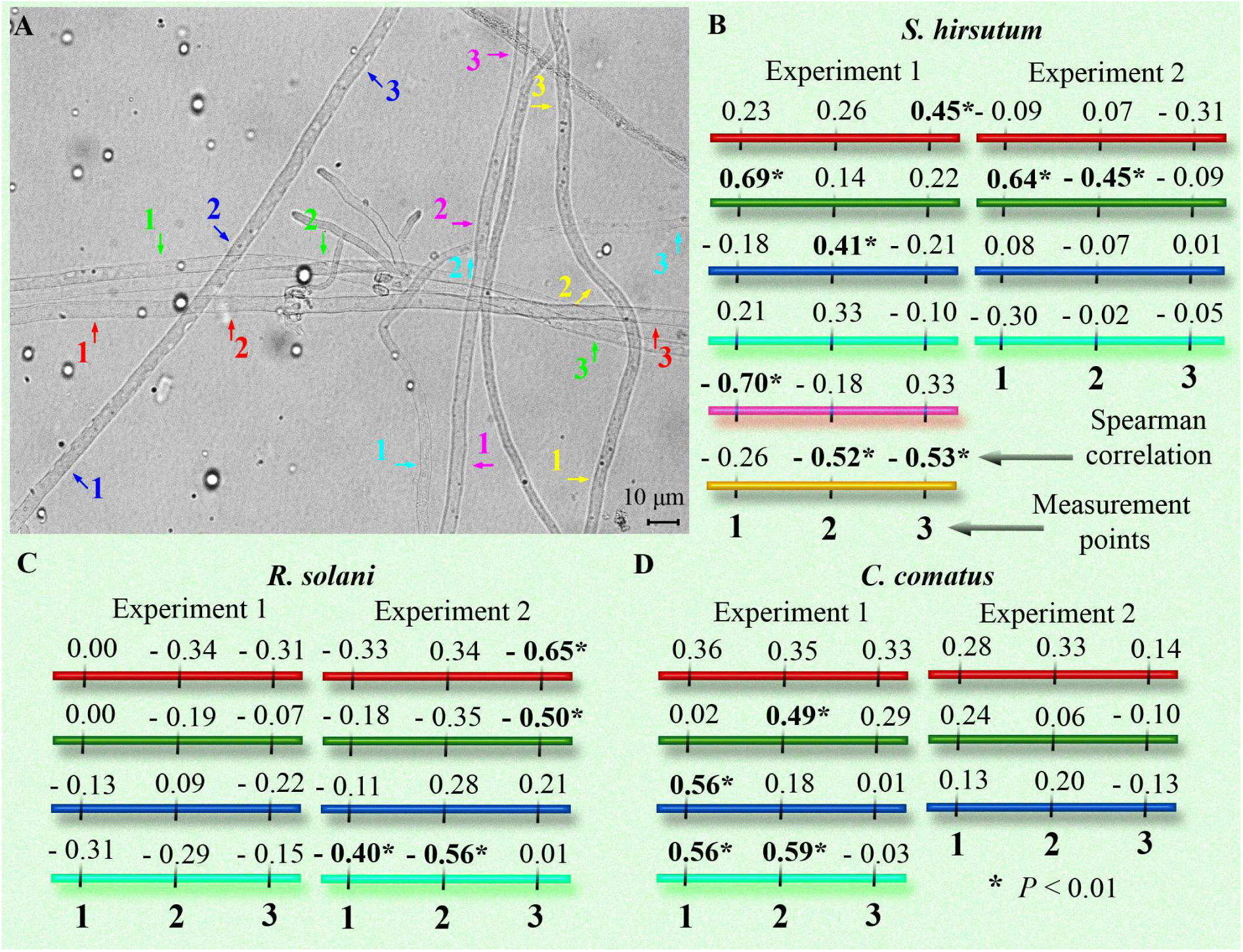
The dynamic experiments. A – frame from a time-lapse video of Experiment 1 of *S. hirsutum*. Three measurement points are marked on each analyzed hypha. The marks on each hypha have their own color; in the diagram on the right (B), the color corresponds to the schematic representation of the hyphae. B–D: schematic representation of the analyzed hyphae. The numbers above each hypha correspond to the Spearman coefficient (r_s_) between the hypha diameter at the measurement point and time (measurements were taken every 60 s for an hour). The dependence is considered significant at |r_s_| ≥ 0.40 and *P* < 0.01 (highlighted in bold and with an asterisk, respectively). If r_s_ is negative, it indicates a gradual shrinkage of the hypha section; if positive, it indicates expansion. If |r_s_| < 0.40, there is either no significant change in diameter at the point, or the diameter fluctuates without a general tendency toward shrinkage or expansion.

## 4. DISCUSSION

### 4.1. Rapid change in hyphal diameter as a universal mechanism for preserving the integrity and functionality of fungal cells

As mentioned in the Introduction, many yeast fungi, some filamentous ascomycetes, and a few species of zygomycetes and basidiomycetes exhibit very rapid cell shrinkage when placed in hyperosmotic conditions (Morris et al. 1986; Slaninova et al. 2000; Lew, Nasserifar 2009; Atilgan et al. 2015; Ene et al. 2015; Mazheika et al. 2020b; Couttenier et al. 2022; Chevalier et al. 2023, 2024). Most often, the fungal cell shrinks as a whole – the protoplast remains adhered to the inner side of the cell wall, and only in some cases is plasmolysis observed. In yeast, the cell wall can thicken instantly and significantly (more than just from physical compression of the stretched cell wall; Slaninova et al. 2000; Ene et al. 2015). It is believed that the modulus of elasticity of the fungal cell wall is close to isotropic, meaning the elastic properties of the wall are the same in both the longitudinal and transverse directions of the elongated cell (Atilgan et al. 2015; Municio-Diaz et al. 2022). Therefore, when the volume of an elongated fungal cell changes, its radius will change faster than its length. This has been confirmed empirically; for example, in *S. pombe*, which has short rod-shaped cells, the shrinkage in length will be half as much as the shrinkage of the transverse diameter (Atilgan et al. 2015). In filamentous fungi, longitudinal shrinkage or extension of hyphae can be largely neglected, as most hyphae adhere to the substrate, creating additional resistance to such changes. This adherence has little effect on changes in hyphal diameter. Therefore, the diameter (radius) of the hyphae serves as the main indicator of changes in the volume of the fungal cell and the elastic strain of the cell wall.

This study shows that three different species of filamentous basidiomycetes, belonging to various systematic and ecological-trophic groups with different average hyphal diameters and growth rates, are subject to rapid changes in hyphal diameters when there is a sharp change in osmolarity and other environmental conditions. This result suggests that the instantaneous shrinkage or expansion of hyphae or yeast cells is a universal structural-protective mechanism characteristic of many fungi. This mechanism allows the fungus to maintain the viability of its cells and quickly restore their functionality under abrupt changes in external conditions (stresses), as well as to maintain continuous functionality of cells under minor or gradual changes in conditions.

The mechanism of shrinkage and expansion of fungal cells is a process determined not only by the laws of physics but also controlled by the cell through the combined action of various cellular structures. In our work on the curtain model of fungal cell organization (Mazheika, Kamzolkina 2024 preprint), four main cellular components were identified as being involved in this mechanism: the elastic cell wall, strong adhesion of the plasma membrane to the cell wall, macroinvaginations of the plasma membrane, and the actin cytoskeleton. With rapid, minor, or slow changes in external conditions and uneven distribution of conditions along the hypha’s length, the actin cytoskeleton complements or replaces the action of turgor pressure, providing tension to the plasma membrane. The plasma membrane is tightly adhered to the cell wall but remains mobile relative to the inner layer of the cell wall. By forming, complicating, increasing in size, and disbanding macroinvaginations, the plasma membrane quickly and sensitively responds to changes in cell size and the shrinkage or expansion of the elastic cell wall. As a result, the fungal cell finely and quickly adapts to the osmolarity and other parameters of the substrate without needing rapid regulation of turgor pressure in the cell. By changing its size, maintaining tension (due to actin and adhesion), and adjusting the functional surface (due to macroinvaginations) of the plasma membrane, the cell can continue vital functions without pausing for rehabilitation.

In the case of abrupt and strong changes in external conditions, such as exposure to 0.6 M sorbitol, the curtain model cannot describe structural processes in the fungal cell. This is primarily because actin cables, an important component of the curtain model, are temporarily depolymerized under stress conditions (Slaninova et al. 2000; Walker et al. 2006; Bitsikas et al. 2011; Elhasi and Blomberg 2019). However, the remaining components continue to perform the structural-protective function. The cell wall shrinks or expands following the protoplast, preventing damage and, in some cases, instantly thickens. The plasma membrane continues to form or disband the macroinvaginations necessary to maintain its integrity. In critical cases, the fungus begins to form larger macroinvaginations, especially pronounced in xylotrophic basidiomycetes, which can form macroinvaginations in the form of thick tubes more than a micron thick and several tens of microns long (Mazheika et al. 2022).

In this study, three important results were obtained that complement and expand the current model and are crucial for understanding fungal physiology. (i) It was found that the studied basidiomycetes do not have a general tendency to rapidly restore their initial size, and therefore turgor pressure, after exposure to an influencing factor like sorbitol. In this aspect, they differ from *N. crassa* and yeast, which begin to increase turgor after osmotic shock within a few dozen minutes and almost restore their initial size and turgor pressure after about an hour (Lew and Nasserifar 2009; Ene et al. 2015). In this regard, basidiomycetes are somewhat similar to oomycetes. This strategy allows basidiomycetes to achieve great physiological plasticity. For example, in mycorrhizal or xylotrophic basidiomycetes, which in soil or dead wood undergo constant changes in general and local conditions, such a strategy allows for the rapid adjustment of turgor functions by the actin cytoskeleton when changes are weak and may be unevenly distributed. In addition, in case of strong changes, they can continue life activity with any average value of turgor pressure, postponing turgor regulation until external conditions stabilize.

(ii) It has been shown that F-actin depolymerization can affect hyphal diameter. However, this effect applies only to S. hirsutum and may result in no change, slight shrinkage, or slight expansion of the average hyphal diameter. There is no dose dependence within the concentration range of 50-200 µM. The dose independence can be explained by the fact that a concentration of 50 µM, especially for LatA, is already quite high and likely leads to significant destruction of F-actin structures. The effect of actin depolymerizers on the diameter of hyphae is crucial for developing the curtain model. This result indicates that actin cables not only regulate the tension of the plasma membrane, as shown earlier (Mazheika et al. 2020b), but also affect the size and shape of the fungal cell. Actin resists turgor pressure, maintains the elastic cell wall, and equalizes the thickness of the hypha along its length, as pressure may vary in different parts of the hypha. The form-restraining and turgor-resisting functions of actin explain why actin depolymerization leads to varying results in different experiments. In mycelium mounted on a microscopic slide, where there is a general tendency for hyphae to expand or shrink due to small osmotic differences and which is restrained by the actin cytoskeleton, the average diameter changes after treatment with inhibitors. Mycelium without such a tendency is not affected by actin destruction. The varying effects of actin assembly inhibitors on different fungal species are likely due to the distinct expression of the form-restraining function of actin in each fungus. In contrast, the membrane tension function may be more universally expressed.

Mechanical stress can affect the diameter of hyphae through its impact on actin. When cells are mechanically damaged, actin cytoskeleton elements may rapidly disassemble, disrupting the form-restraining function. However, the slight shrinkage of *S. hirsutum* hyphae under mechanical stress can be attributed to damage to the plasma membrane and a technical error due to deterioration in the quality of the preparation.

(iii) Another important result is that plasmolysis in moderate hyperosmotic conditions in basidiomycetes can occur, but it is specific. In one of our previous works, it was shown that in *R. solani*, plasmolysis in 0.6 M sorbitol, as in many fungi, is rare (Mazheika et al. 2020b). However, in that work, there was an incubation period between sorbitol treatment and observation, which did not allow the detection of rapid reversible concave plasmolysis. This phenomenon occurs in the first minutes after placing the sample in a hyperosmotic medium and quickly disappears. Rapid reversible concave plasmolysis serves as a backup mechanism for macroinvaginations of the plasma membrane. When there is a sharp drop in turgor pressure, the macroinvagination system fails to manage its task and cannot pack excess membrane into invaginations in time. Concave plasmolysis compensates by removing the excess membrane in large folds, which remain attached to the cell wall at points of strongest focal adhesion. In this state, the cell retains its integrity but is unlikely to function fully. Therefore, after 5–10 min, plasmolysis disappears and is replaced by macroinvaginations if necessary.

Why can rapid reversible plasmolysis without preliminary incubation in liquid medium be observed in hyperphase only in *R. solani*, but not in the other two fungi? On the one hand, as will be shown below, *R. solani* most likely has a slightly lower average turgor than *S. hirsutum* and *C. comatus*. This may explain why the same concentration of sorbitol has different effects on the fungi studied. On the other hand, the differences in turgor pressure are unlikely to be so great as to react so differently to sorbitol. It can be assumed that basidiomycetes from different ecological-trophic groups use structural-protective mechanisms differently. In xylotrophs (and possibly in other basidiomycetes such as mycorrhizal, copro-humus, and others), the system of macroinvaginations is more developed and complex than in the endophyte/phytoparasite *R. solani*. Therefore, under certain conditions, with a sharp change in the osmolarity of the environment, fungi with a complex system of macroinvaginations cope with the shock as the plasma membrane quickly packs or unpacks into macroinvaginations. *R. solani* resorts to plasmolysis since its macroinvagination system is not designed to handle such abrupt changes in pressure and cell size.

The question arises: why does sorbitol cause rapid reversible concave plasmolysis in *S. hirsutum* after an hour of preincubation in liquid malt medium, similar to that observed in *R. solani* without preincubation? It is necessary to consistently analyze the stages of experiments with the preincubation of *S. hirsutum* mycelium. Preincubation, as experiments demonstrate, can cause a slight expansion of hyphae in *S. hirsutum*. Our previous work showed that preincubation also causes an increase in the number of lomasomes (the main type of macroinvaginations in xylotrophs; Mazheika et al. 2022). The apparent contradiction (when the hyphae expand, the lomasomes should disband) can be explained as follows: the osmolarity of the cytoplasm of the mycelium growing on a dense nutrient medium is, on average, slightly higher than the osmolarity of the mycelium growing in a liquid medium of the same composition as the dense medium. When preparing microscopic specimens with liquid malt medium, a partial imitation of surface growth occurs due to the thinness of the liquid layer on the slide and other factors. Therefore, the tendency for hyphae to expand is not observed. Possibly, for the same reason, water as a mounting liquid does not cause mycelium expansion. An hour-long incubation in excess liquid medium results in a slight expansion of the hyphae and an increase in turgor. Moreover, after an hour, similar to the case with a hypertonic medium, the mycelium does not attempt to restore turgor to its original state. De novo membrane synthesis also begins as a response to cell expansion and increased turgor. In an hour, the equilibrium between synthesis and expansion is not achieved, and the excess membranes are packed into macroinvaginations. In the same work (Mazheika et al. 2022), it was shown that a 12-hour preincubation does not lead to an increase in the number of lomasomes; an equilibrium state is already established within this period. Then the preincubated mycelium is placed in 0.6 M sorbitol. Since the turgor pressure is slightly higher than during preincubation, the shrinkage of the hyphae occurs to a lesser extent. However, there is an excess of plasma membrane in the cells, packed into macroinvaginations. There is a need to form macroinvaginations and increase the size and complexity of the existing ones. There is likely a permissible limit to the number of macroinvaginations in terms of membrane surface area. The checkpoint system is triggered, and permission for concave plasmolysis is initiated as a safety mechanism when the macroinvagination system reaches its capacity limit.

### 4.2. Physiological significance of the influence of permeabilizing agents on changes in hyphal diameter and plasmolysis

Non-ionic detergent Triton X-100 and the proteolytic enzyme Proteinase K are traditionally used in biology for cell permeabilization, which increases the permeability of the plasma membrane to improve the penetration of antibodies and various probes into the cell. Triton X-100 is known to disrupt the integrity of the bilipid layer of the membrane and protein-lipid bonds but has little effect on protein conformation (Johnson 2013). Proteinase K, on the contrary, causes permeabilization by destroying transmembrane proteins or connections between membrane proteins, weakly affecting the bilipid layer itself (Amidzadeh et al. 2014).

Usually, the cell wall of a living fungal cell is under elastic strain. The elastic strain directly depends on the forces or pressures acting on the wall and inversely depends on the thickness of the cell wall and the elastic modulus (an indicator of the wall’s elasticity, depending on its molecular composition, structure, cross-linking of its polysaccharide fibrils, etc.; Atilgan et al. 2015; Municio-Diaz et al. 2022). This strain can be estimated by the change in hyphal diameter relative to the diameter of relaxation. The diameter of relaxation is the diameter of the hypha when no forces or pressures (such as turgor, actin resistance or its contractile force) act on the cell wall (Atilgan et al. 2015; Mazheika, Kamzolkina 2024 preprint). To estimate elastic strain and turgor, the deflation assay is often used. In this case, the cell membrane is microperforated with a laser. If the fungal cell was under pressure, it decreases in volume to a state close to relaxation of the cell wall. However, since the adhesion of the protoplast to the cell wall remains, it continues to affect the elastic strain of the wall (Atilgan et al. 2015; Davi et al. 2019; Chevalier et al. 2023, 2024). The treatment of mycelium with Triton X-100 in the current study is somewhat analogous to the deflation assay. In the permeabilized protoplast, the turgor pressure drops to zero. Most likely, the actin cytoskeleton is also destroyed, but even if it continues to resist, this is not so important since in many cells convex plasmolysis occurs – the adhesion of the plasma membrane to the cell wall is disrupted. Detachment of the protoplast from the cell wall allows for complete relaxation of the cell wall. Triton X-100 treatment could be used instead of deflation assays to measure turgor and elastic strain of the wall as a more accurate method. However, the method requires improvement since not all mycelial cells undergo plasmolysis, resulting in inaccurate results. Additionally, it is unknown how Triton X-100 affects the cell wall itself and its elastic properties.

In our experiments, if to consider the hyphal diameters in Triton X-100 as approximate diameters of cell wall relaxation, it turns out that 0.6 M sorbitol reduces the turgor pressure in the studied fungi to nearly zero on average, as the shrinkage values in sorbitol and Triton X-100 are close and do not differ statistically. However, in *R. solani*, the overall shrinkage in sorbitol exceeds the shrinkage in Triton X-100 by a few percent, which may indicate that the average turgor pressure in this fungus is slightly lower than in the other two fungi. 0.6 M sorbitol compresses the hyphae of *R. solani* below the relaxation point of the cell wall – the protoplast, shrinking, drags the cell wall with it, leading to compression of the cell wall itself.

Another interesting question concerns the reasons for strong convex plasmolysis in fungal hyphae in Triton X-100. Such plasmolysis indicates not only protoplast permeabilization and a drop in turgor but also a significant disruption of the adhesion of the cell membrane to the wall, even in areas of strong focal adhesion that remain during concave plasmolysis. Adhesion of the plasma membrane to the cell wall is thought to be mediated by both single molecules and focal sites (clusters of molecules concentrated in specific sites in the membrane). Both focal and single molecule adhesion may be mediated by mechanosensory proteins that are transmembrane inserted into the plasma membrane and anchored in the cell wall by their sensory domains. Another possibility is GPI-APs or GPI-glucans, which are anchored by their ceramide tails in the plasma membrane, with the protein or glucan portion embedded in the cell wall (Elhasi, Blomberg, 2019; Komath et al., 2022; Mazheika, Kamzolkina 2024 preprint). Since Triton X-100 weakly denatures proteins but actively destroys lipid layers, it can be assumed that GPI compounds play a major role in the adhesion of the plasma membrane to the cell wall in fungi. On the other hand, Triton X-100 can disrupt adhesion by destroying lipid-protein bonds and releasing mechanosensors from the membrane. Similarly, the weaker effect of Proteinase K on the probability of plasmolysis supports the hypothesis of GPI adhesion versus mechanosensory adhesion. Moreover, it is possible that different types of adhesion dominate in different cases. For example, in thin hyphae of *C. comatus*, convex plasmolysis occurs after treatment with Proteinase K, which may indicate the dominance of protein adhesion. On the other hand, Proteinase K has a much weaker effect on hyphal diameter, indicating its weak and species-specific permeabilizing properties, as well as possibly suboptimal conditions for proteolysis (it is known that Proteinase K is more active at elevated temperatures and in the presence of detergents). In this way, the nature of fungal adhesion of the plasma membrane to the cell wall requires further research.

## 5. CONCLUSION

Unlike most land plant cells, which are protected by integumentary, protective, and mechanical tissues from external conditions, fungal cells of substrate hyphae are in direct contact with the substrate. To a certain extent, fungi can resist changes occurring in the substrate through the synthesis of hydrophobins, the formation of external capsules or matrices, melanization of the cell wall, changes in membrane permeability, etc. However, these protective mechanisms and structures are not always sufficient for rapid adjustment to minor fluctuations in external conditions, and they cannot preserve the active living hypha’s functionality during sudden and strong changes. Therefore, fungi took a different path: rather than swimming against the current, they allow themselves to be carried by it until reaching a quiet backwater. Mechanisms for preserving the functionality of fungal cells include the ability of fungi to quickly change the diameter of their hyphae in response to various stress conditions and minor fluctuations, primarily changes in the osmolarity of the liquid phase of the substrate. At the same time, the fungal cell maintains its integrity and the tension of the plasma membrane, and quickly restores metabolic activity. Additionally, fungi such as basidiomycetes do not waste resources on rapid restoration of turgor pressure, adjusting cell size and other parameters to the current pressure. This mechanism employs elasticity and other properties of the cell wall, turgor regulation mechanisms, plasma membrane macroinvagination systems, adhesion of the plasma membrane to the cell wall, and the actin cable system. However, our knowledge about the named physiological processes and structures that are of fundamental importance in the life of fungi remains ephemeral. Therefore, extensive, high-quality microscopic studies analyzing the dynamics of fluorescently labeled actin cables in basidiomycetes; searching for moleculas that ensure adhesion of the plasma membrane to the cell wall and studying connections between them and actin; analyze the dynamics of the distribution of turgor pressure and tension of the plasma membrane along the fungal hypha and other studies are needed.

## Supporting information

Caption for Supplementary Video

