## Supplementary material for "Instantaneous change in hyphal diameter in basidiomycete fungi": Caption for Supplementary Video

**LEGEND FOR SUPPLEMENTARY VIDEO**

**Video**. Time-lapse video obtained in an experiment of dynamic changes in *R. solani*. The microscopic slide is mounted with malt medium containing 0.6 M sorbitol. The same field of view was captured at 1-min intervals for an hour. Concave plasmolysis occurs in some cells during the first 8 minutes, then disappears, leaving only a small convex plasmolysis near individual septa. After 20 minutes, it disappears as well. Scale bar 20 µm.
